# Wideband ratiometric measurement of tonic and phasic dopamine release in the striatum

**DOI:** 10.1101/2024.10.17.618918

**Authors:** Amy Gottschalk, Haley Menees, Celine Bogner, Semele Zewde, Joanna Jibin, Asma Gamam, Dylan Flink, Meea Mosissa, Faith Bonneson, Hibo Wehelie, Yanaira Alonso-Caraballo, Arif A. Hamid

## Abstract

Reward learning, cognition, and motivation are supported by changes in neurotransmitter levels across multiple timescales. Current measurement technologies for various neuromodulators (such as dopamine and serotonin) do not bridge timescales of fluctuations, limiting the ability to define the behavioral significance, regulation, and relationship between fast (phasic) and slow (tonic) dynamics. To help resolve longstanding debates about the behavioral significance of dopamine across timescales, we developed a novel quantification strategy, augmenting extensively used carbon-fiber Fast Scan Cyclic Voltammetry (FSCV). We iteratively engineered the FSCV scan sequence to rapidly modify electrode sensitivity within a sampling window and applied ratiometric analysis for wideband dopamine measurement. This allowed us to selectively eliminate artifacts unrelated to electrochemical detection (i.e., baseline drift), overcoming previous limitations that precluded wideband dopamine detection from milliseconds to hours. We extensively characterize this approach *in vitro*, validate performance *in vivo* with simultaneous microdialysis, and deploy this technique to measure wideband dopamine changes across striatal regions under pharmacological, optogenetic, and behavioral manipulations. We demonstrate that our approach can extend to additional analytes, including serotonin and pH, providing a robust platform to assess the contributions of multi-timescale neuromodulator fluctuations to cognition, learning, and motivation.

## Introduction

A central feature of substance abuse and psychiatric illness is the breakdown of dopamine (DA) regulation across brain regions and timescales ^1–3^. Extensive evidence indicates that striatal DA release *can* fluctuate on fast “phasic” and slow “tonic” timescales to regulate adaptive behavioral and cognitive flexibility. Yet, the behavioral significance and mechanistic basis of these changes are poorly understood, largely due to technical limitations in bridging timescales of DA measurement. For example, sustained DA tone is hypothesized to regulate the energization and maintenance of motivated behaviors from seconds to hours, depending on task complexity ^4–6^. This view is extensively supported by microdialysis measurements, the gold standard for calibrated DA measurement over minutes and hours. These multi-minute DA elevations are observed during heightened arousal and vigor, such as those produced by psychoactive drugs or task epochs with elevated reward rates and motivation ^7–9^. However, microdialysis quantifies DA levels in dialysates that contain small fractions of actual brain DA concentrations, requiring several minutes of collection for each sample to accumulate sufficient DA above detection thresholds ^10,11^. Therefore, this slow-sampling bandwidth renders microdialysis unsuitable for capturing rapid, sub-second DA changes, but paradoxically, this very property has been utilized in the literature to define the notion of a slow-varying “tonic” DA level in the striatum, without specification of the relevant frequency band ^6,12,13^.

By contrast, transient, sub-second DA release in the striatum is tightly linked to the burst-firing of DA cells responding to unpredicted rewards or cues predictive of future rewards ^14,15^. These surprise-driven “phasic” DA events support an influential Reinforcement Learning (RL) theory that emphasizes DA’s role in learning by signaling reward prediction errors (RPEs) ^16,17^. In addition to extensive electrophysiological spike recording in the midbrain, this view is supported by DA concentration ([DA]) measurements in the striatum using electrochemical, fast scan cyclic voltammetry (FSCV) ^9,18–20^ and genetically-encoded fluorescent DA sensors ^21–25^. However, these fast, sub-second DA measurement approaches are unstable on long timescales due to non-linear changes in fluorophore expression/bleaching or background drifts in FSCV, narrowing their bandwidth and precluding their use for comparing DA changes across minutes or hours.

Informed by these quantification methods with non-overlapping bandwidths, details about the multi-timescale contributions of DA to flexible behavioral control, interactions between tonic and phasic DA signals, and circuit regulators of DA across regions and timescales remain intensely debated across theoretical rifts. For example, one perspective posits that DA supports learning and motivation on *separate* timescales ^5,13^, with fast phasic signals relaying RPEs for learning and ambient tonic levels regulating motivation. By contrast, other frameworks suggest that timescales of DA variation dynamically interact in service of behavioral flexibility ^6,26,27^. For example, we recently proposed that in the ventral striatum (a region consistently linked to microdialysis “tonic” DA changes correlated with motivational vigor), fast DA fluctuations relay discounted reward expectation across task-relevant timescales ^9^. Because surprise is proportional to expectation and, in turn, updates prediction, this view posits a bidirectional inverse relationship between fast and slow timescales of DA releases ^6,26^. While these frameworks are conceptually attractive, resolving the details of how DA RPEs used for learning accumulate to set ambient DA tone that shapes performance remains elusive due to a lack of DA measurement approaches that effectively bridge behaviorally relevant timescales.

Here, we present a novel wideband quantification strategy that builds on the FSCV technique by employing ratiometric principles to overcome technical challenges in broadband DA measurement. By using asymmetric adsorption durations to rapidly modulate electrode sensitivity to electroactive analytes within a single sampling, we achieved selective amplification of DA redox signals while effectively eliminating baseline drift artifacts. After extensive validation of this approach both *in vitro* and *in vivo* with simultaneous microdialysis, we demonstrated its utility by assessing how phasic [DA] fluctuations consistently accumulate to set tonic levels under reward, optogenetic, and pharmacological conditions. Our method is generalizable to additional electroactive brain analytes, including serotonin, and holds promise for potential applications in human intracranial measurements. Ultimately, this technical advancement provides a powerful platform for rigorous testing of theoretical predictions about how broadband neuromodulator fluctuations support the computational demands of behavioral flexibility across timescales.

## Results

### FSCV adsorption epoch controls electrode sensitivity to analytes

We sought to devise a strategy for quantifying wideband DA across physiologically relevant timescales, utilizing the extensively validated electrochemical FSCV method ^28–30^. Standard FSCV repeatedly applies a triangular voltage sweep (typically −0.4V to 1.3V) onto a carbon-fiber microelectrode to promote a brief redox reaction between the carbon fiber surface and nearby electroactive analytes ^31^ (**Fig. 1A**). After amplification and digitization, the detected current during the voltage sweep is proportional to the concentration of analytes, with different molecules undergoing redox at different potentials. Parameters of the voltage ramp have been extensively iterated for selectivity and sensitivity to various analytes, including DA ^30^. Importantly, in the absence of analytes, the carbon fiber generates a substantial capacitance current that must be subtracted from all samples ^32^ (**Fig. 1B**). This background subtraction uncovers the current contribution of analytes under study, a redox signature trace known as the voltammogram (**Fig. 1B**, *pink trace*). In the context of *in vivo* recordings, this background subtraction step is typically repeated every 20-30 seconds (as in ^9,33,34^) because the large capacitance current is non-stationary due to microscopic changes at the surface of the carbon fiber ^35,36^. As such, FSCV is a *differential* method requiring iterative background correction, restricting its application for comparing measurements across minutes and hours.

**Figure 1:**
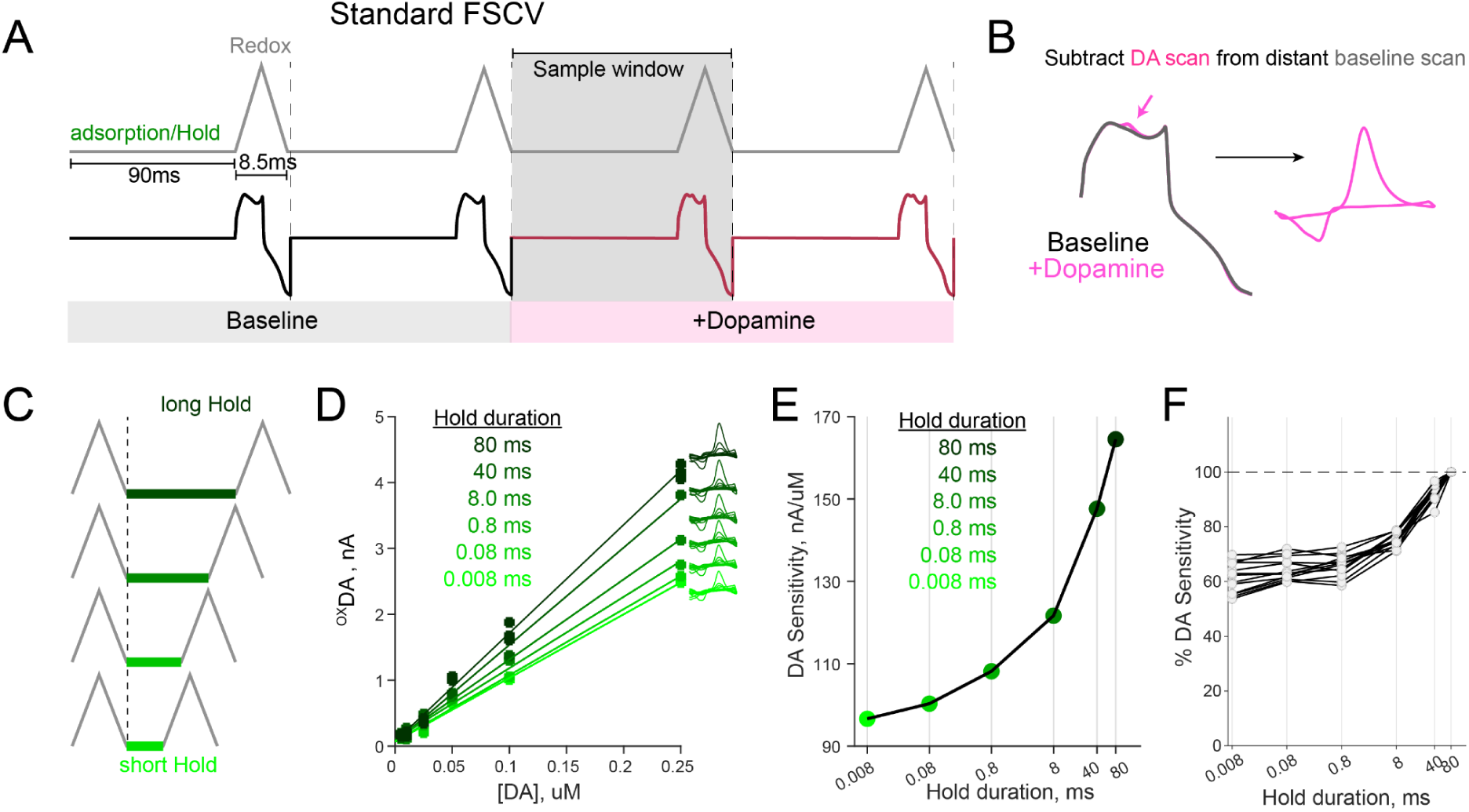
FSCV Hold duration modulates sensitivity. **(A)** Illustration of standard FSCV scan sequence. (*Top*) Short triangular voltage ramps are applied following a prolonged adsorption (Hold) period in each sampling window (gray-shaded region). (*Bottom*) Example Faradaic capacitive current traces in the absence (black) and presence (red) of DA. **(B)** A background capacitive current from a distant (30s) time point is typically subtracted at each sample to reveal the analyte-specific current. Note that because the capacitive current changes over time, the experimenter has to decide the optimal temporal gap for background separation. **(C)** Illustration of changing the Hold/adsorption period to assess how it affects the sensitivity of measurements. Decreasing the Hold period results in faster sampling rates. **(D)** Calibration curves of the same electrode to repeated [DA] exposure under varying Hold durations. **(E)** The effect of Hold duration on the calibration slope or sensitivity (same electrode as shown in D). **(F)** The percent change from maximal sensitivity due to changing Hold duration across all electrodes (each gray line is an electrode).

We reasoned that the background correction problem might be resolved using ratiometric detection methods inspired by their application in noise canceling ^37^, sensor self-calibration ^38^, and repetition suppression in fMRI ^39^. In principle, ratiometric detection relies on signal quantification with two (or more) measurements/sensors that have differential sensitivity to eliminate various noise sources ^40^. To translate this method to FSCV, we explored whether the sensitivity of the carbon fiber to DA can be modified without affecting the electrochemical backbone of the FSCV method. We focused on the inter-voltage-sweep-interval (here, interchangeably referred to as “Adsorption” or “Hold” period) as a candidate modulator of electrode sensitivity. This Hold epoch precedes the triangular ramp and is typically held at a negative potential to facilitate the electrostatic attraction of positively charged DA molecules to the electrode ^36,41^.

We therefore sought to test if decreasing this Hold period under standard FSCV would systematically change the sensitivity of the carbon fiber to [DA] in a flow cell preparation (**Fig. 1C**). We compared the magnitude of DA oxidation current (^OX^DA) as we applied a range of [DA] while varying the Hold duration from 8 microseconds to 80 milliseconds. Each electrode was tested on the full range of [DA] and Hold durations to isolate their effect on the sensitivity of the same carbon-fiber sensor. We found that shortening the Hold duration significantly decreased electrodes’ sensitivity across a range of [DA] (**Fig. 1D**, One-way rmANOVA; main effect of Hold duration on sensitivity: F(6,130) = 277.2, p =1.5 x 10^−71^). While the relationship between sensitivity and Hold duration was non-linear (**Fig. 1E**), we observed, on average, a 38% reduction in calibration slope by reducing the hold duration by a factor of 10,000, from 80 milliseconds to 8 microseconds across electrodes tested (**Fig. 1F**).

### vhFSCV for ratiometric quantification of DA concentration

Armed with this insight, we next designed a scan sequence that would facilitate the comparison of currents for pairs of Hold durations within a single sampling period. Instead of using physically separated probes with different sensitivities, our strategy relied on rapidly modifying a single electrode’s sensitivity to enable ratiometric quantification at the same brain microdomain (**Fig. 2A**). Specifically, we introduce the variable-hold FSCV (vhFSCV) with a 100ms sampling period, and each sample constituted two triangular scans with parametrically varying Hold periods.

**Figure 2:**
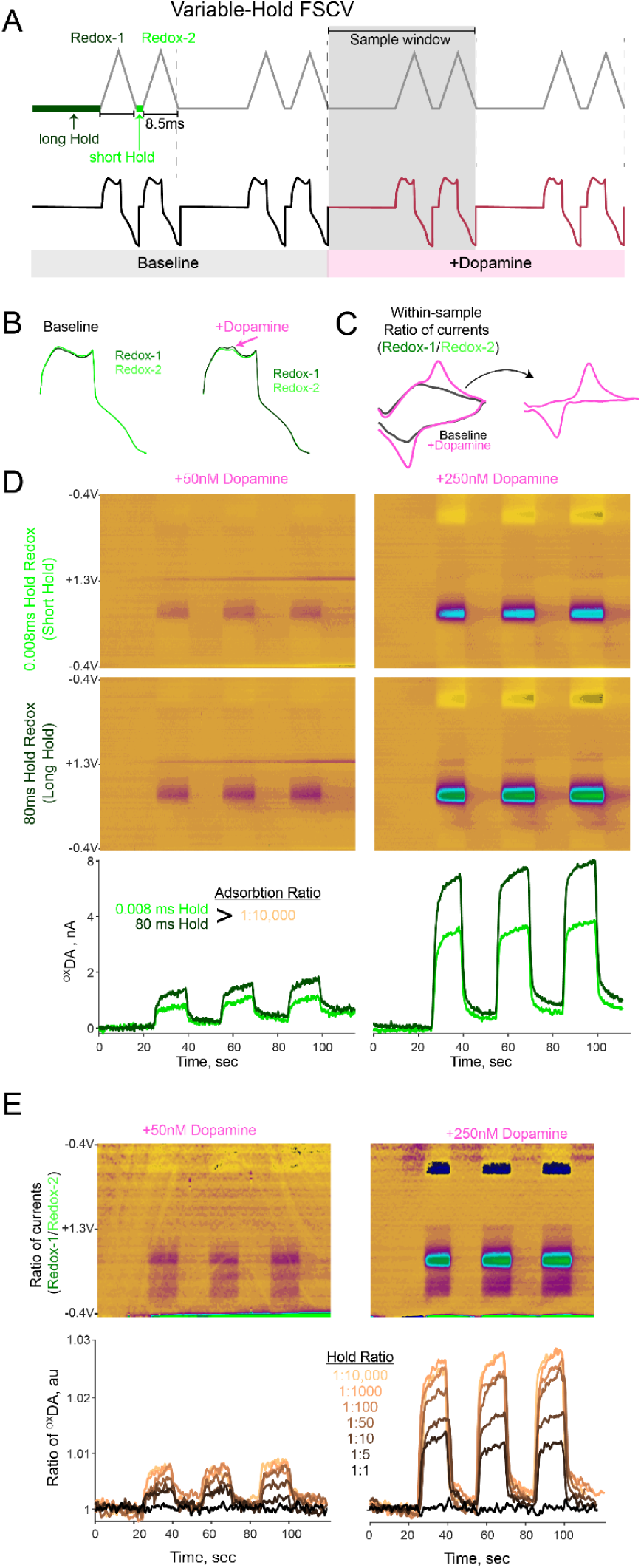
VhFSCV allows for ratiometric measurement of DA redox and background currents for within-sample baseline subtraction. **(A)** (*Top*) Depiction of vhFSCV scan sequence wherein a second triangular voltage ramp is added within a sample window (gray), and the preceding adsorption epoch length is parametrically varied for each scan, resulting in a scan with long adsorption (Redox-1) and a scan with short adsorption (Redox-2). (*Bottom*) Example of corresponding faradaic current trace for vhFSCV in the absence and presence of DA. **(B)** During the baseline (non-DA) periods, capacitance currents from Redox-1 and Redox-2 are very similar. When DA is applied, the faradaic currents remain similar but diverge in current contributions from DA (pink arrow). **(C)** Computing a ratio of currents detected at each potential performs a within-sample normalization of capacitive currents. The ratio of currents for the baseline epoch had a distinct shape, and this ratio during DA administration was specifically divergent at DA oxidation potentials. Thus, subtracting these curves retrieves that canonical DA voltammogram. **(D)** (*Top)* Color plots of the short-Hold and long-Hold plots when administering 50nM and 250nM DA in a flow cell. The top and bottom currents are acquired in rapid succession within-sample (see A), but separated here to demonstrate that the DA current magnitudes from the two scans are different. *(Bottom)* Current magnitudes at the peak DA oxidation potential, normalized to the start, for a 0.008ms Hold and 80ms Hold successive scans (Hold Ratio of 1:10,000). **(E)** *(Top)* Color plots of the ratio of currents at each potential within a sampling period for a 1:10,000 Hold ratio. Note that this ratio color plot facilitates the inspection of redox signatures of DA or other analytes differentially detected due to Hold asymmetry. *(Bottom)* Traces of the ratio of currents, specifically at the peak oxidation of DA, for several combinations of Hold ratios for the same [DA] administered.

We hypothesized that when the Hold durations for Scan-1 and Scan-2 were identical, the contribution of the capacitance and analyte redox currents would be identical. Moreover, with progressively asymmetric Hold durations across the two scans, we posit that the analyte-specific redox currents would differ proportional to the Hold duration asymmetry (reflecting their modulated sensitivity). Critically, currents unrelated to analytes undergoing redox (such as faradaic currents) would be *similar* for the two scans. In other words, an asymmetric Hold epoch would separate detected currents into those differentially dependent on the electrochemistry promoted by the Hold duration. Consequently, this strategy has the advantage of within-sample normalization of the faradic current that primarily contributes to electrode drifts across long timescales (**Fig. 2B**) without affecting the properties of the redox reaction (**Fig. 2C**).

We tested these predictions in the flow cell by applying various [DA] onto an electrode and separately analyzing either the currents detected (^OX^DA) for each triangular scan, or the ratio of oxidation currents across the scans (^OX^DA-Ratio) in each sample. We found that in response to the 50nM or 250nM [DA], scans with short Hold durations (0.008ms) reported a smaller ^OX^DA current than scans with longer Hold durations (80ms) (**Fig. 2D**).

Because these differences are within a single sampling window, this difference in sensitivity indicated that ^OX^DA-Ratio may be proportional to absolute [DA]. To test this possibility, we first calculate the “Hold-Ratio” of multiple combinations of adsorption epochs at each scan (e.g., *Scan1-vs-Scan2*: 40ms-vs-40ms = 1:1 ratio, 8ms-vs-80ms = 1:10 ratio, 0.008ms-vs-80ms = 1:10,000 ratio), and asked whether parametrically varying this Hold-Ratio predicts proportional differences in the ^OX^DA-Ratio currents across the two scans (**Fig. 2E**). Indeed, for the 50nM and 250nM [DA] tested, progressively increasing the Hold-Ratio (stronger asymmetry in adsorption periods) enhanced the difference in detected DA across the two scans, yielding higher ^OX^DA-Ratio (**Fig. 2E**). Note that because the Current-Ratio is calculated for each sample, at all voltage potentials of the triangular ramp, we can observe the current signatures (as in **Fig 2E**, *top*) and inspect the voltammogram of analytes undergoing differential redox due to the asymmetric Hold duration. Finally, when the Hold-Ratio was 1 (i.e., equal adsorption periods), the two Scans reported identical ^OX^DA currents, resulting in a constant ^OX^DA-Ratio of 1 during DA application (**Fig. 2E**, *black trace*), supporting our negative-control prediction. These results indicate that our modified two-scan approach allows for within-sample ratiometric quantification of DA changes.

### The ^OX^DA-Ratio is linearly proportional to DA concentration across timescales

Next, we characterized the changes in electrode sensitivity that give rise to this ratiometric property, constraints on analyte species, and stability over timescales. We exposed eight electrodes to [DA] that span physiological levels and assessed how ^OX^DA and ^OX^DA-Ratio change. Across the tested [DA] range, we observed that symmetric Hold durations (Hold-Ratio = 1:1) had identical calibration curves (**Fig. 3A**, *left*). Progressively increasing the Hold-Ratio across the two scans significantly separated the slopes of calibration curves between short and long adsorption scans (**Fig. 3A**, One-way rmANOVA; main effect of Hold-Ratio on slope difference: F(6,42) = 122.8, p =5.2 x 10^−25^). Reflecting this incremental separation in sensitivity across Hold-Ratios, we observed that the ^OX^DA-Ratio for each DA run increased for increases of both Hold-Ratio and [DA] (**Fig. 3B**) in 8 electrodes tested (**Fig. 3C**, 2-way rmANOVA; main effect of [DA]: F(1, 7) = 138.4, p = 2.3 x 10^−20^, main effect of Hold-Ratio: F(1, 7) = 51, p = 4.3 x 10^−18^). Moreover, the effect of Hold-Ratio on ^OX^DA-Ratio is linear for all electrodes tested (all p > 0.25, Ramsy RESET test for non-linearity). Finally, this divergence in sensitivity is selective to DA, as demonstrated by a significant correlation between the voltammograms derived from the Current-Ratio and the individual scans (**Fig. 3D, E**). These results lead us to conclude that our ratiometric-FSCV approach quantifies the ratio of DA oxidation currents (^OX^DA-Ratio) that linearly covary with DA concentration.

**Figure 3:**
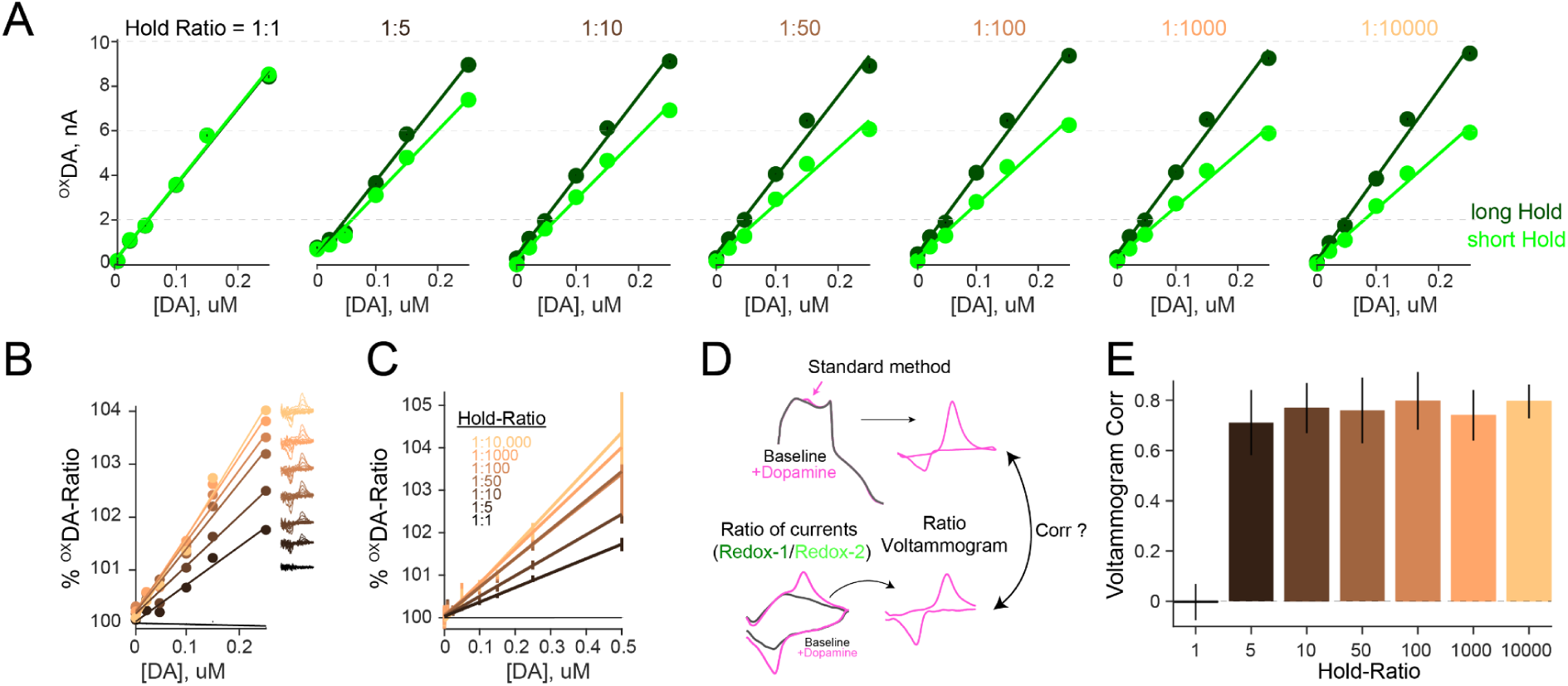
Proportional change in sensitivity across Hold-Ratios underlies ^OX^DA-Ratio quantification of DA concentrations. **(A)** DA calibration curves for short and long Hold durations across Hold-Ratios for an example electrode. For each Hold asymmetry, darker colors indicate the response of the more sensitive scan (longer Hold), and lighter colors indicate currents of the less sensitive scan (shorter Hold). Note the progressive deviation of the calibration slopes with progressively asymmetric Hold-Ratios. **(B)** This sensitivity divergence can be captured as the ratio of currents detected at the peak oxidation current (^OX^DA-Ratio) linearly proportional to administered DA. Note that smaller hold ratios still provide a proportional quantification of [DA] but with less contrast. The right inset indicates voltammograms for DA redox currents across Hold-Ratios. **(C)** The same plot as (B) shows the average data across eight electrodes tested. **(D)** To assess if voltammograms for the individual scans were similar to voltammograms for the ratiometric method, we compare their correlations. **(E)** Correlations between voltammograms for ratiometric detection and Scan-1 under the standard method demonstrate a high correlation between signatures for DA redox currents.

By measuring the ratio of currents detected between two scans, our observations support the hypothesis that vhFSCV is linear and selective for analytes that are differentially detected by the two scans due to Hold duration asymmetry. Consequently, we predict two desirable outcomes. First, this logic should hold for additional analytes typically investigated by FSCV, such as pH ^42^ or serotonin ^43,44^, as they also rely on electrostatic attraction during the Hold duration to facilitate FSCV detection. Second, the exclusivity of the Current-Ratio to differentially detected analytes renders vhFSCV insensitive to signal contributions that do not depend on the Hold-Ratio asymmetry, such as baseline drift caused by faradic current changes. This is crucial for wideband quantification because it allows the ^OX^DA-Ratio to compare “phasic” and “tonic” [DA] fluctuations across timescales without contamination from non-biological signal sources.

To test the first prediction, we extended our flow cell characterization to varying serotonin concentrations [5-HT], or phosphate-buffered saline (PBS) acidified at different pH units. Supporting our predictions, we find the same patterns of results for modulation of Current-Ratio to graded [5-HT] and pH levels compared to the DA experiments. Specifically, when Hold durations are identical for the two scans, the ratio of oxidation currents for serotonin(^OX^5-HT-Ratio) or pH (^OX^pH-Ratio) did not vary across different concentrations (**Fig. 4A, B**; *black circles*). Moreover, progressively increasing the Hold-Ratio or analyte concentration increased ^OX^5-HT-Ratio (**Fig. 4A**, 2-way ANOVA; main effect of Hold-Ratio: F(6, 30) = 7.8, p = 4 x 10^−5^) and ^OX^pH-Ratio (**Fig. 4A**, 2-way ANOVA; main effect of Hold-Ratio: F(6, 30) = 5.6, p = 4 x 10^−4^) linearly (all p > 0.25, Ramsy RESET test for non-linearity). These results indicate that our ratiometric principles are extendable to the measurement of serotonin and pH. Moving forward, we focus on the maximal Hold-Ratio achieved across datasets (1: 400 or 1:10,000) that provide the largest sensitivity difference across all analytes and concentrations tested.

**Figure 4:**
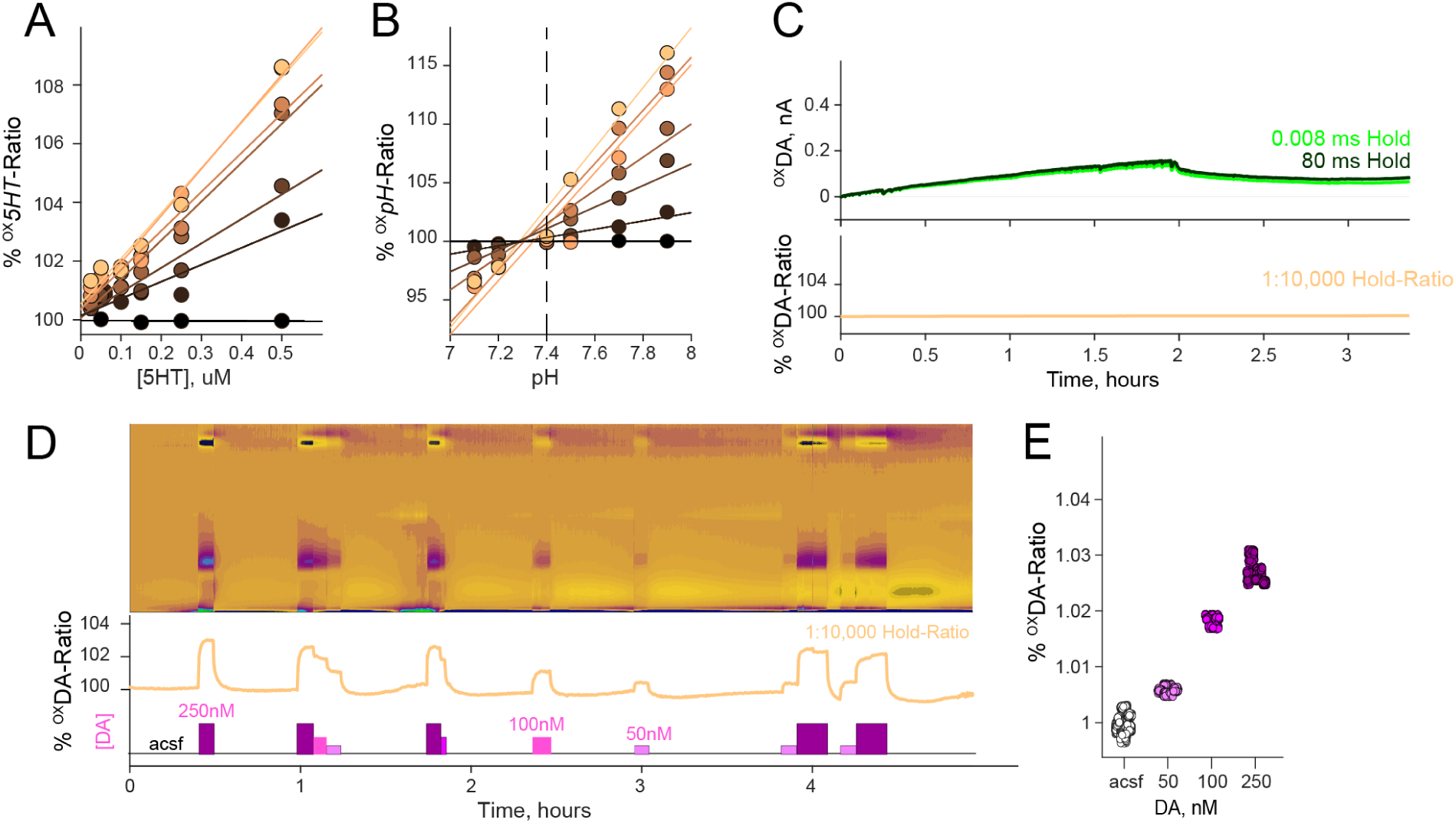
Constraints on analyte species and stability over timescales of vhFSCV. **(A)** Same as Figure 3B. We tested the effect of varying serotonin concentrations [5-HT] and Hold-Ratio on the ratio of detected currents at the serotonin oxidation potential (^OX^5HT-Ratio). **(B)** Same as (A) for the effect of pH and Hold-Ratio on ^OX^pH-Ratio. **(C)** Currents at the DA oxidation potential for short (green) and long (black) hold scans and the Ratio-Current at a Hold-Ratio of 1:10,000 (yellow) of an electrode in aCSF for 3.5 hours. Note that while the currents for the individual scans dynamically drift, the Current-Ratio remains stable over hours. **(D)** Electrode exposed to 5-minute pulses of varying [DA] across 5 hours at Hold-Ratio 1:10,000. **(E)** Quantification of ^OX^DA-Ratio for aCSF and various [DA] across the 5-hour recording session demonstrating sensitivity to DA is preserved across hours.

We next tested the second prediction that vhFSCV is insensitive to signal contributions that do not depend on the Hold-Ratio, allowing wideband [DA] quantification from milliseconds to hours. We examined whether the ^OX^DA-Ratio changed over several hours without any electroactive analytes present, with electrodes submerged in PBS for hours. While the ^OX^DA currents of both scans unpredictably drifted as previously reported ^35^, these drifts were proportional to each other, leading to a stable ^OX^DA-Ratio over hours (**Fig 4C**).

To determine if the sensitivity to [DA] is preserved across this extended window, we exposed the electrode to five-minute-long steps of different [DA] in a separate 5-hour-long recording session to synthetically mimic [DA] tone changes observed *in vivo*. We found that the ^OX^DA-Ratio was stable in PBS response over hours and proportionally increased in magnitude for [DA] separated by hours (**Fig. 4F**; one-way rmANOVA, main effect of [DA]: F(3,15) = 377.5, p = 2.4×10^−14^). These results lead us to conclude that vhFSCV can detect wideband DA changes across seconds to hours in the flow cell preparation and turned to *in vivo* characterization.

### Validation of vhFSCV [DA] measurement with simultaneous microdialysis in vivo

Pharmacological antagonism of the D2 autoreceptor and the dopamine reuptake transporter (DAT) has been extensively documented to change phasic and tonic [DA] fluctuations in the striatum ^8,33^. We leveraged these established findings as a testbed for assessing whether vhFSCV accurately quantifies striatal [DA] changes across timescales, validated with simultaneous microdialysis in awake, head-restrained mice. Specifically, we implanted a microdialysis guide cannula directed at the right Nucleus Accumbens (NAc) and a second burr hole targeting the same region for acute vhFSCV (**Fig. 5A**). We calibrated the microdialysis HPLC-ED with [DA] standards (**Fig. 5B**), determined the probe’s recovery rate and delay (**Fig. 5C**) to quantify striatal [DA] in five-minute intervals (see methods for details). We then lowered the microdialysis probe and vhFSCV electrode into the NAc, separated by 300-500 microns. After stabilization, we simultaneously recorded [DA] in NAc using vhFSCV and microdialysis for an initial baseline period and injected 10mg/kg i.p. of the DAT antagonist, nomifensine.

**Figure 5:**
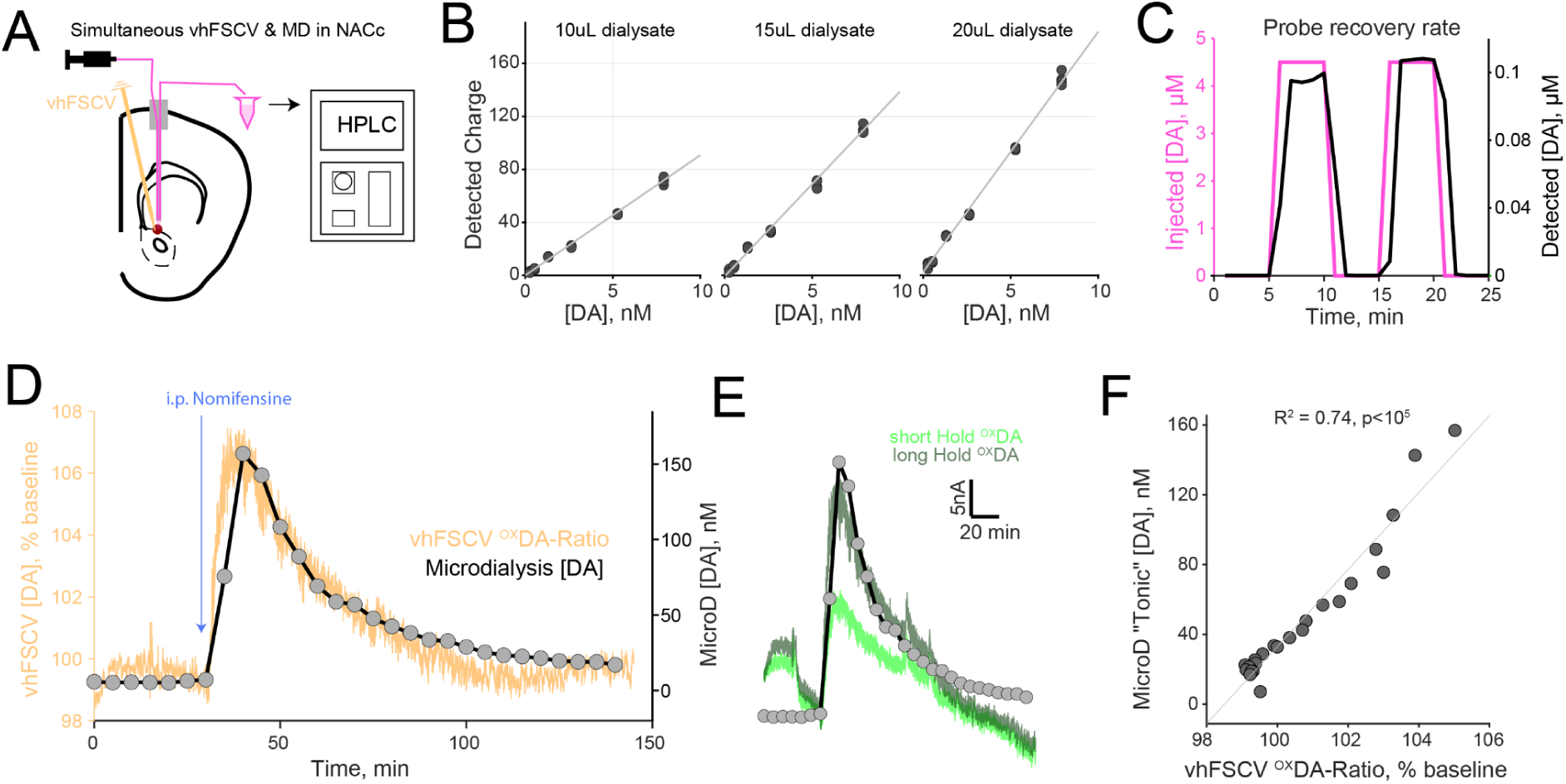
Validation of vhFSCV wideband DA measurement in vivo with simultaneous microdialysis. **(A)** Simultaneous measurements of wideband NAc DA using vhFSCV and microdialysis. We targeted a microdialysis probe and carbon fiber to the same striatal microdomain 300-500 μm apart. **(B)** Microdialysis HPLC-ED calibration curves for [DA] standards across different dialysate volumes before the start of the *in vivo* experiment. **(C)** Probe recovery fraction (amount of [DA] recovered from known [DA]) and collection delay were determined for each microdialysis probe before recording sessions. **(D)** Microdialysis [DA] (gray) aligned with simultaneous vhFSCV ^OX^DA-Ratio (orange) during baseline levels and i.p. injection of 10mg/kg nomifensine. Note the strong similarity in the time course of the two time series. **(E)** Same as (D) for the timecourse of microdialysis [DA] re-aligned with ^OX^DA of individual short-Hold (light green) and long-Hold (dark green) scans. Note common drift-like deviations early in the session normalized by the ratio measured in (D). Moreover, nomifensine-induced DA increase is differentially detected by the two scans in proportion to the modulated sensitivity. **(F)** Correlation between microdialysis [DA] measurements and vhFSCV ^OX^DA-Ratio (R^2^ = 0.74, p<10^5^).

We observed phasic [DA] fluctuations as quantified by ^OX^DA-Ratio signal during the baseline period, and the administration of nomifensine significantly elevated the ambient DA tone for several minutes accompanied by large [DA] transients (**Fig. 5D**). Moreover, simultaneous microdialysis [DA] quantified at 5-minute intervals was better correlated with the timecourse of ^OX^DA-Ratio (R^2^= 0.74), compared with ^OX^DA of the long- or short-Hold scans (R^2^= 0.63 and R^2^= 0.49 respectively) which shared drift-like deviations that were normalized by the ratiometric division (**Fig. 5E, F**). This close match between ^OX^DA-Ratio and microdialysis [DA] levels suggests that vhFSCV quantifies broadband tonic and phasic [DA] dynamics *in vivo*.

### Deploying vhFSCV to survey wideband [DA] fluctuations across striatal regions and experimental manipulations

The behavioral significance, mechanistic basis, and relationship of multi-timescale DA changes are poorly understood. In this section, we demonstrate the utility of vhFSCV in advancing a deeper understanding of how phasic [DA] fluctuations affect tonic levels. To this end, we recorded wideband [DA] changes under behavioral, pharmacological, and optogenetic manipulations of subsecond DA transients to assess whether phasic events consistently accumulate to shape DA tone.

In the first experiment, we used pharmacological blockade of the D2 autoreceptor by systemic injections of raclopride followed by nomifensine. We found that this schedule of drug administration elevated phasic DA events in the NAc for several minutes with distinct patterns of tonic [DA] change (**Fig. 6A**). Specifically, raclopride significantly increased both the frequency and amplitude of phasic transients. In contrast, the blockade of DAT with nomifensine produced even larger phasic transients that had broader half-widths (**Fig. 6B**) together with a large increase in tonic [DA] levels. We found a similar effect of nomifensine on transient [DA] in multi-site vhFSCV recordings in the medial and lateral dorsal striatum (DMS or DLS, **Fig. 6C, D**).

**Figure 6:**
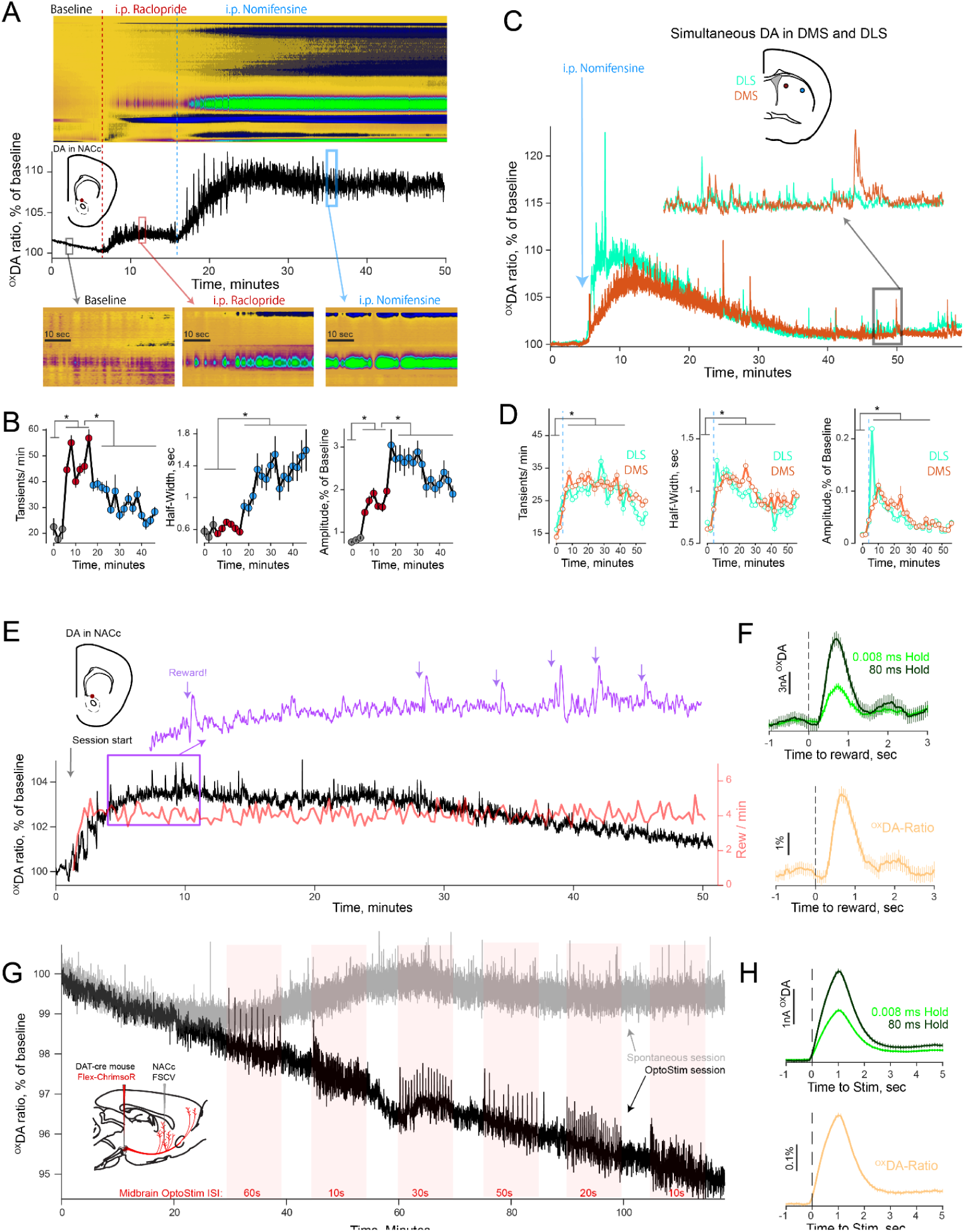
Quantification of wideband [DA] changes across striatal regions following pharmacological, reward, and optogenetic manipulations using vhFSCV. **(A)** Color plot of Current-Ratio (*Top)* and ^OX^DA-Ratio (*Middle)* of an *in vivo* vhFSCV (Hold-Ratio 1:10,000) recording in the NAc while administering i.p. injections of 1 mg/kg raclopride and 10 mg/kg nomifensine. Note the step-like increase in DA tone across pharmacological conditions that are of similar magnitude to the largest phasic increases observed at the onset of nomifensine. (*Bottom)* Representative Current-Ratio color plots to indicate vhFSCV phasic transients during the baseline, raclopride, and nomifensine epochs. **(B)** Quantification of changes in DA transient frequency, half-width, and amplitudes across drug conditions. Stars indicated significant differences between phasic statistics (two-sample Kolmogrov-Smirnov test, p<0.05). **(C)** Simultaneous, two-channel vhFSCV in medial and lateral dorsal striatum with systemic administration of nomifensine. **(D)** Same as (B) for quantification of transient statistics in dorsal striatum. **(E)** NAc vhFSCV during reward expectation task (black), and the reward rate per minute (red). Mice received rewards every 8-15 seconds, with progress to rewards indicated by audiovisual stimuli. Note the overall increase in DA tone as the mouse receives rewards. Purple inset indicates phasic reward responses at the end of each trial. **(F)** (*Top*) Alignment of ^OX^DA-Ratio to reward delivery for the short- and long-Hold scans. Note that the more sensitive, long-Hold scans report larger current deflections than their less sensitive counterparts to the same reward-evoked DA release event. (*Bottom*) Alignment of the current ratio. **(G)** NAc vhFSC in NAc during optogenetic stimulation of the midbrain. After a baseline epoch, mice received bursts of midbrain stimulations with indicated inter-stimulation-intervals for 10 minutes (red-shaded region). Note the overall decline in DA tone quantified by vhFSCV in opto stim session (black) but not in duration-matched spontaneous sessions (gray). **(H)** Same as (F) for alignment of optogenetic stimulation.

In a second experiment, we recorded NAc DA changes in a mouse trained on the Pavlovian cue-reward association task previously demonstrated ^24^ to drive reward anticipation and DA release in the striatum (**Fig. 6E**). In this task, auditory tones escalated in frequency to indicate progress to a delayed reward delivery (4-8 sec) on a trial-by-trial basis. We noted phasic [DA] increases at reward delivery that were differentially detected by vhFSCV scans with asymmetric Hold durations (**Fig. 6F**). Moreover, the wideband ^OX^DA-Ratio timecourse strongly correlated with the reward rate (R^2^ = 0.17) as previously demonstrated ^9^, indicating that our quantification approach is sensitive to *task-relevant* tonic and phasic [DA] changes.

Finally, in a third experiment, we expressed a cre-dependent, red-shifted excitatory opsin, ChrimsonR selectively in dopaminergic cells in DAT-cre mice and quantified broadband [DA] changes during optogenetic stimulations (**Fig. 6G**). Specifically, we stimulated midbrain DA cells with repeated 1s long 630nm light pulses at 60Hz in blocks of 10 minutes (each with different inter-stim intervals) followed by unstimulated 5-minute blocks. While each midbrain stimulation led to phasic increases in [DA] (**Fig. 6H**), we observed a striking and persistent decrease in [DA] tone as quantified by the ^OX^DA-Ratio across all six sessions tested (**Fig. 6G**, *black trace*), in contrast to duration-matched spontaneous sessions without stimulation (**Fig. 6G**, *grey trace*). These results suggested that phasic DA increases correlate with elevated [DA] tone under specific conditions, prompting us to characterize this relationship in additional detail.

### A paradoxical constraint on the accumulation of phasic [DA] transients into [DA] tone

Prevailing computational models in RL suggest that phasic DA increases encode RPEs that accumulate to set tonic [DA] levels correlated with reward rates ^5,27,45^. However, the specifics of this accumulation process remain contested. Some frameworks invoke passive mechanisms, where the frequency of phasic DA RPEs determines the power in tonic bandwidths ^13,23,46^. In contrast, another perspective posits that active intrastriatal circuits may prospectively transform phasic [DA] into tonic [DA] ^6,26^. Such mechanisms may allow for a rapid and flexible adjustment in ambient DA levels based on cues or internal states, extending the effective bandwidth of task-relevant DA “tone” to include multi-second ramps ^9,47^. Floresco and colleagues report that tonic and phasic [DA] may be independent communication channels regulated by orthogonal midbrain mechanisms ^48^. These contrasting possibilities remain unresolved because the specific frequency band(s) that specify striatal tonic [DA] is elusive. The sample bandwidth of microdialysis (>5-10 mins) has achieved an *implicit consensus:* the idea that synaptic spillover of phasic [DA] is detected as DA tone ^12,49^ is attractive from a quantification ^50,51^ and modeling perspective ^46,52–54^. However, the dependence of tonic [DA] on the frequency and amplitude of phasic transients has not been concretely tested *in vivo*.

Leveraging the wideband vhFSCV dataset summarized above, we performed an initial analysis to clarify whether phasic transients accumulate into tonic levels. We first asked which ^OX^DA-Ratio frequency bands correlate with the [DA] quantified by microdialysis in our dual vhFSCV-microdialysis recordings (**Fig. 5D**). Using spectral analysis methods, we decomposed the wideband ^OX^DA-Ratio into its time-frequency representation (**Fig. 7A**), and correlated the subsampled power to the microdialysis data. We found that, as expected, frequency bands less than 0.003Hz (>5 min period of microdialysis) had strong correlations (R^2^ > 0.55, p<0.01), suggesting that sample collection in microdialysis imposes a low-pass filter of DA fluctuations (**Fig. 7B**). Interestingly, additional frequency bands centered near 0.1Hz (10-15s period) also exhibited even greater correlations with microdialysis levels (R^2^ =0.8, p<0.0001; **Fig. 7B**), indicating that multisecond fluctuations typically described as ramps may significantly contribute to microdialysis levels detected. These results are not affected by potential synchronization jitter between vhFSCV and microdialysis.

**Figure 7.**
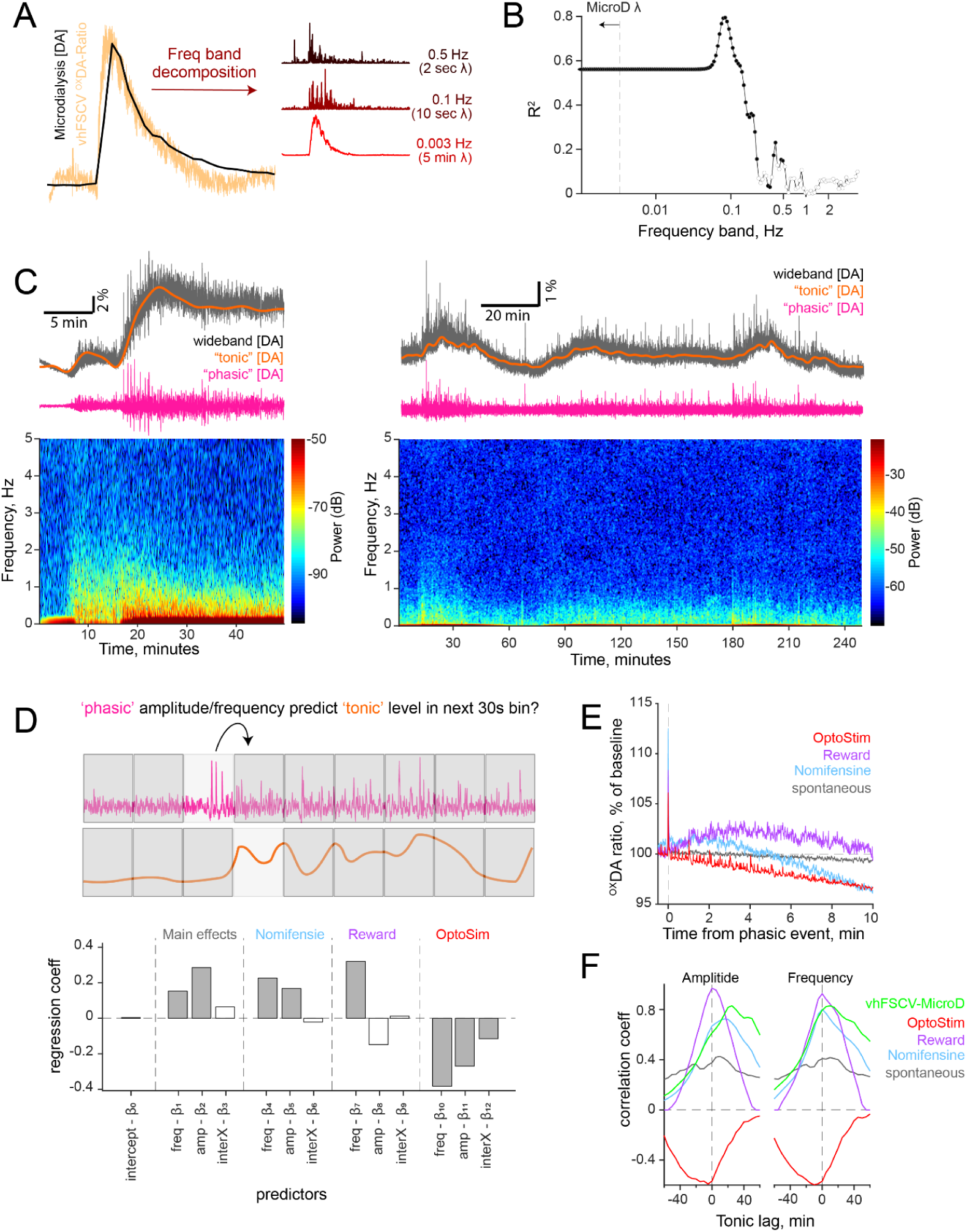
Assessing whether phasic DA events consistently accumulate to set DA tone. **(A)** Illustration of frequency band decomposition of vhFSCV. Dual recording session shown in 5D. **(B)** For each vhFSCV frequency band, we computed its Pearson’s correlation to microdialysis results and reported the coefficient of determination. Significant correlations (p<0.05) are indicated with black circles. **(C)** (*Top*) We operationally defined “tonic” DA in vhFSCV as changes in frequency bands highly correlated to microdialysis samples in Figure 6B. Thus, we lowpass filtered the wideband OXDA-Ratio to derive “tonic” DA fluctuations (orange). By comparison, fluctuations faster than 0.5Hz can be defined as “phasic” changes (pink). (*Bottom*) Spectrogram of wideband vhFSCV OXDA-Ratio. Note shifts in frequency power across the recording window. (*Left*) Pharmacology session is the same as 6A. (*Right*) Spontaneous sessions without external stimuli or manipulation. **(D)** (*Top*) Illustration of analysis. We compared how phasic amplitude and frequency in wideband vhFSCV OXDA-Ratio, averaged in 30s windows, predicted “tonic” levels computed as in (C) in the subsequent 30s window. (*Bottom*) Results of a mixed-effects regression model demonstrating strong regression coefficients for phasic frequency and amplitudes in predicting tonic levels across conditions. Filled gray bars represent significant regressors, and hollow bars are not. **(E)** Phasic-event triggered alignment of wideband vhFSCV across a 10-minute window for various conditions. **(F)** Cross-correlation analysis of variables derived in (D) assessing the maximal temporal offset of “tonic” DA to maximize correlations with the mean amplitude (*Left*) and frequency (*Right*) of phasic events in 30-second bins.

We next asked whether phasic ^OX^DA-Ratio event rate or amplitudes predict [DA] tone from 25 sessions (7 mice) under spontaneous, nomifensine, reward, and optogenetic manipulations. Tonic [DA] was operationally defined as fluctuations occurring over periods slower than 5 minutes (i.e., <0.003 Hz), in line with the microdialysis bandwidth from above. We used a low-pass filter with a cutoff of 0.003 Hz to compute vhFSCV “tonic” [DA] (**Fig. 7C**), detected phasic events, and assessed how transient rate and amplitude predicted subsequent [DA] tone in sliding 30s analysis windows (**Fig. 7D**). We constructed a hierarchical mixed-effects model to predict tonic [DA] using phasic event amplitude, frequency, and their interaction as predictors, with experimental conditions as fixed effects and random intercepts for animals and sessions. We found significant main effects of phasic amplitude (β_amp_ = 0.29, p = 4 ×10^−32^) and frequency (β_freq_ = 0.15, p = 7 ×10^−11^) on [DA] tone in subsequent 30-second windows, suggesting that, under spontaneous conditions, phasic [DA] events contribute to the accumulation of tonic [DA]. A similar pattern of results was observed during dopamine reuptake blockade with nomifensine (β_amp_ = 0.11, p = 1 ×10^−6^, β_freq_ = 0.22, p = 5 ×10^−11^), consistent with the hypothesis that phasic transients can pool in the extracellular space to elevate tonic dopamine levels.

Interestingly, during the reward expectation task, we observed a selective significant effect of transient frequency on predicting DA tone (β_freq_ = 0.32, p = 4 ×10^−4^), while phasic amplitude was not significantly predictive, though trending in the negative direction (β_amp_ = −0.15, p = 0.15; **Fig. 7D**). These results hint at the possibility that frequent reward-evoked transients (encoding RPEs) are more likely to promote elevated tonic DA than large amplitude deflections.

The strong positive coefficients under spontaneous, drug, and reward conditions led us to predict that optogenetically driving phasic [DA] would similarly increase DA tone. Surprisingly, we found that exogenous phasic events promoted a decreasing DA tone with significantly negative regression coefficients for both frequency (β_freq_ = −0.38, p = 0.04) and amplitude (β_amp_ = −0.27, p = 0.04; **Fig. 7D**). These results show that artificial phasic [DA] events do not accumulate into DA tone, hinting at additional circuit (or pharmacodynamic) interactions that may regulate the incorporation of phasic DA events into DA tone beyond passive synaptic overflow. These overall findings are also supported by transient-triggered wideband [DA] alignment, where we observe significant DA tone following phasic events across datasets, while optogenetically induced phasic events decreased successive DA levels (**Fig. 7E**). Moreover, cross-correlation analyses with sliding tonic DA relative to bins of phasic DA events indicate a rightward shift in correlations. These results suggest that phasic events generally lead to elevated DA tone several minutes after, while opto-stimulation produced a negative relationship between DA tone and phasic transient amplitude/frequency (**Fig. 7F**).

## Discussion

Here, we introduce vhFSCV, a novel quantification method designed to bridge the gap between rapid phasic and sustained tonic DA measurements. Traditional approaches have restricted bandwidths, constraining a comprehensive understanding of DA’s role in reward learning, cognition, and motivation. To overcome these limitations, we redesigned the measurement scan sequence in FSCV and leveraged ratiometric quantification principles. Our strategy rapidly modulated electrode sensitivity through asymmetric Hold durations across successive electrochemical scans within a sampling period. This change in sensitivity likely arises from modulated electrostatic adsorption at the negatively charged electrode surface ^41,55^. We found that systematically modulating electrode sensitivity scaled the detected redox currents in proportion to both analyte concentration and Hold duration asymmetry. We exploited these proportionalities to derive a ratiometric measure, ^OX^Ratio, which we demonstrate effectively isolates analyte-specific signals from noise artifacts independent of the adsorption duration. This performance benefit was sustained for several hours in the flow-cell preparation.

We further validated these characteristics in the striatum of awake mice undergoing behavioral, pharmacological, and optogenetic manipulations. We first performed simultaneous recording of vhFSCV and microdialysis in the NAc as mice received the DA reuptake antagonist, nomifensine, to elevate tonic and phasic DA. Our analysis revealed that ^OX^Ratio is tightly correlated with microdialysis DA measurements, and this ratio signal normalized various noise and drift artifacts shared by the two scans. Moreover, we applied spectral analysis to the wideband ^OX^Ratio signal and identified at least two frequency band regimes that strongly correlated with microdialysis [DA] fluctuations. These included bands near the five-minute sampling period of microdialysis (∼0.003Hz), and another near the 0.1Hz period of DA ramps ^9,47,56^. These results provide clarity to a longstanding ambiguity about the relevant frequency bands of “tonic” DA changes in previous microdialysis reports.

We additionally demonstrate the utility of our measurement approach in assessing longstanding debates on the temporal relationship between phasic and tonic DA fluctuations. Phasic DA elevations are associated with surprising environmental feedback and are hypothesized to drive Value learning in RL frameworks by signaling RPEs. These updated Value signals drive higher reward expectation, motivational engagement, and strongly correlate with tonic DA levels in the striatum. However, details about whether phasic transients accumulate to set ambient DA tone, and in turn, if high DA tone constrains the magnitude of reward-evoked phasic release, are central to current theoretical debates in RL ^4–6,13,26,27,57,58^. We leverage vhFSCV to provide an initial assessment of whether phasic DA transients consistently predict the elevation of tonic DA levels under various conditions. Our analysis revealed that under spontaneous and pharmacological blockade of DA reuptake, the magnitude and frequency of phasic DA transients predict DA tone in wideband vhFSC measurement. These findings are consistent with early “synaptic spillover” hypotheses ^12,49^. This interpretation was partially supported in a reward-learning task; only the frequency, but not magnitude, of phasic events predicted DA tone increases. Finally, artificially introducing phasic DA release through brief optogenetic stimulation did not causally elevate the tonic vhFSCV frequency bands. This preliminary assessment reveals a paradoxical constraint on the accumulation of phasic DA into tonic levels, providing an initial basis for future investigations into circuit mechanisms that may shape this interaction under different task demands.

The ratiometric approach in vhFSCV is parameter-free and does not require modeling changes in the Faraday current over time. Moreover, while the implementation of vhFSCV here uses a 10Hz sampling rate, the ratiometric advantages we characterize here extend to faster (or slower) sampling rates, provided that the Hold duration asymmetry is preserved. This contrasts with recent electrochemical detection efforts to bridge timescales of DA quantification. Modifications to the postprocessing pipeline ^59^, the adsorption epoch ^60–62^, and the scan waveform ^63–66^ in FSCV have provided robust methods to study tonic DA *in vivo* over hours ^43,64,65,67,68^. However, these methods have restricted temporal resolution, limiting the ability to measure phasic transients that occur on the timescale of milliseconds. Additional efforts to mitigate this limitation have used techniques to estimate capacitive background currents to remove electrode drift ^69–73^. While promising, modeling capacitive background currents has proved difficult as it requires strong assumptions and carefully fabricating electrode micro surfaces.

Although we focused our ratiometric analysis on raw oxidation currents of analytes under study, one limitation of our approach is that we did not implement advanced chemometric analyses with dimensionality reduction, which has been demonstrated to facilitate robust multivariate calibration ^74–76^. Indeed, these procedures can be trivially implemented in the data analysis pipeline because the Current-Ratio is evaluated sample-wise in vhFSCV. Moreover, vhFSCV does not modify the electrochemical backbone of standard FSCV; as such, the full suite of typical postprocessing pipelines developed for FSCV can readily migrate to vhFSCV.

In sum, we present vhFSCV as a powerful tool for accurately quantifying broadband neurotransmitter dynamics *in vivo* that will facilitate a deeper understanding of computational and mechanistic principles supporting behavioral and cognitive flexibility across timescales. This feature is grounded in the observation that ^OX^Ratio in vhFSCV is selectively proportional to redox current sources sensitive to the Hold asymmetry we imposed. Thus, it logically follows that our ratiometric quantification is a readout of the absolute neurotransmitter levels undergoing redox. Consequently, we argue here that our ratiometric approach not only normalizes noise artifacts but, if calibrated appropriately, offers a quantitative assessment of absolute neurotransmitter concentrations across timescales.

## Acknowledgements

We thank Michael Frank, Joshua Berke, Eric Newman, Janet Dubinsky, Hoawei Zhang for feedback on earlier versions of the manuscript, and members of the Hamid Lab for feedback at various stages of the project. We also thank Mitch Roitman for donating FSCV micromanipulators, Paras Patel for sharing FSCV headstages and drawings, Christopher Moore, Sinda Fekir, and Matthew Nassar for early discussions on this quantification approach.

This work was supported by the Howard Hughes Medical Institute (Hanna Gray Fellowship), UMN Dean’s office (ECRA award), National Institute on Drug Abuse (T32DA007234), and National Institute of Neurological Disorders and Stroke (R25NS117356),.

## Author Contributions

AG collected and analyzed the data and contributed to writing the manuscript. HM, and CB performed data collection. SZ, JA, AG, DF, MM, FB, HW and YAC assisted with data collection. AAH conceptualized and developed the method, designed and supervised the study, analyzed the data, and wrote the manuscript.

## Methods

### Animals and Surgery

Adult wild-type C57BL/6J (JAX, 000664) and DAT-cre^+^ (JAX, 020080) mice were used in this study. Mice were kept on a reversed 12:12 light:dark cycle and were tested during the dark phase. All animal procedures were conducted in accordance with the guidelines of the NIH and approved by the University of Minnesota Institutional Animal Care and Use Committee.

We followed standard sterile surgical procedures for all implants and stereotaxic injections of cre-dependent viruses described below. Briefly, mice were anesthetized with isoflurane (5% induction and maintained at 1-2% in 1 L/min oxygen), and body temperature was maintained at 37°C using a heating pad. For all mice, a metal head post was first secured to the skull, and additional implants were inserted according to targeted stereotaxic coordinates. At the end of the experiment, mice were perfused to verify the histological expression of opsin and the accuracy of implant targeting. For *in vivo* vhFSCV experiments, we used either a small burr hole for acute electrode drops or a micromanipulator guide cannula implanted above the right NAc (1.3AP, 0.8ML, relative to bregma), DMS (1ML, 1.0 AP) or DLS (2.5ML, 0AP). An Ag/AgCl reference electrode was cemented in the contralateral cortex. For microdialysis experiments, a guide cannula (Eicom CXG-4, 0.5 mm outer diameter) was implanted above the NAc (1.5AP,1.0ML), with a burr hole drilled into the skull directly adjacent to the guide cannula to target a carbon fiber to NAc (1.5AP; +1.3 ML) at a 15° ML angle during dual vhFSCV and microdialysis experiments.

### Voltammetry

vhFSCV electrode construction was performed as previously described ^9^. Briefly, we used glass-encased carbon-fiber microelectrodes with an exposed carbon fiber length of 150µm - 200µm. The assembled microelectrode was connected to a custom, 4-channel head stage ^77^, and voltage scans, data collection, optogenetic stimulation, reward delivery, and task synchronization were performed using custom LABVIEW software controlling a National Instruments (NI) PCIe-6353 DAQ device.

The vhFSCV scan sequence builds on the standard 8.5ms wide, −0.4V to +1.3V FSCV triangular ramp applied at 100ms intervals. For standard FSCV experiments in Figure 1, we delivered this voltage ramp and parametrically varied the Hold duration, effectively modulating the sampling rate of FSCV. Thus, the hold duration is calculated as the difference between the sampling period and ramp duration. We implemented vhFSCV by delivering two triangular scans within the 100ms sampling period, parametrically varying each scan’s Hold period to achieve the desired Hold-Ratio. Specifically, we created a modified waveform that concatenates i) Scan-1 (8.5ms), ii) Hold duration for Scan-2 (*x*), and iii) Scan-2 (8.5ms) at an 8.5μs (117.53kHz) update rate on the NI DAQ. At the termination of the concatenated waveform, the remaining duration (*y*) serves as the Hold duration of Scan-1 of the next cycle. This strategy allows the user to define a Hold-Ratio (1:1, 1:10, etc), and the software will solve for the duration of *x* based on these two equalities: 100_Sample Period(ms)_ = 8.5_Scan-1(ms)_ + x_(Scan-2 Hold,ms)_ + 8.5_Scan-1(ms)_ + y_(Scan-1 Hold,ms)_ and Hold-Ratio = *y*/*x*. Simultaneous with the writing of the concatenated waveform to an analog output channel connected to the waveform pin on the head stage, we simultaneously read analog signals from DAQ channels connected to working electrodes on the head stage. After truncating input signals at the boundaries of the triangular scans, we stream to disc voltage signals proportional to currents detected at the carbon fiber (100nA/V) using an unlimited FIFO queue to prevent data loss.

All *in vitro,* vhFSCV experiments were performed in a flow cell system ^78^, using a flow-regulated gravity feed system with pneumatic solenoids or linear actuators plunging a syringe with micro stepper motors to inject PBS or DA into the electrode chamber. All solutions were made from 10X stock PBS with a final pH of 7.4. Solutions containing DA and 5HT were prepared immediately before use to avoid oxidative degradation. Known concentrations of DA (1 nM - 1000 nM) and 5HT (5nM - 500nM), and PBS calibrated across a range of pH units (7.1 - 7.9) were tested across Hold-ratios to examine the effects of [DA] and Hold-ratio on ^OX^DA-Ratio. For in vivo vhFSCV, glass-encased carbon-fiber electrodes were acutely lowered into the NAc (−3.3DV, relative to brain surface), DMS (2.5DV), or DLS(2.8DV) using either a stereotaxic apparatus or a micromanipulator in mice implanted with a guide cannula. Once lowered, the electrode was preconditioned before recording by cycling at 60Hz for 15 minutes, followed by an additional 15 minutes at 10Hz, or until stabilized.

### Microdialysis

The recovery rate for a microdialysis probe (Model CX-I-X-Y, diameter 0.22mm, EiCOM) was tested *in vitro* before *in vivo* use by inserting the probe into solutions containing known concentrations of DA. Non-buffered aCSF (composition: 147 mM NaCl, 2.8 mM KCl, 1.2 mM CaCl_2_, 1.2 mM MgCl_2_) was perfused through the probe continuously at a flow rate of 2µl per min. The fraction of DA collected through the probe over 5-minute intervals was averaged and compared against the original [DA] to determine the recovery fraction for each microdialysis probe. To assess tubing delay from DA dialysis to sample collection, we used standard pulse-chase methods where we briefly transferred the probe from aCSF to [DA] and assessed the sample delay to DA detection. The probe was inserted into the NAc, extending 1 mm below the guide cannula, and it was left to equilibrate for at least 60 minutes. A carbon fiber electrode was lowered to the NAc, 300-500μm from the microdialysis probe and was preconditioned as described above. Immediately before starting the *in vivo* session, a 96-well plate was pre-filled with six standard DA calibration samples, and the remaining wells with antioxidant solutions (composition: 0.2 M acetic acid, 2.0 mM oxalic acid, 6.0 mM L-Cysteine at pH 3.2). Dialysate samples were collected once every 5 minutes (10µl total) using a fraction collector (FC-90, AMUZA) and were cooled to 6°C in the dark until the session ended. At the end of a session, a second set of standard DA calibration samples (the same concentration as those added at the beginning of the experiment) were added to the plate to account for any oxidative degradation that may have occurred. High-performance liquid chromatography was performed on dialysis samples using an EiCOM HTEC-600 system equipped with an EiCOM AS-700 autosampler. DA was separated using a reversed-phase separation column (EICOMPAK PP-ODS3) and electrochemically detected using a graphite working electrode with an applied potential of +400mV. Data analysis was performed using Clarity chromatography software to quantify the detected charge at the DA potential in dialysis samples. The recovery fraction and standard DA calibration samples were then used to determine absolute brain [DA], and the tubing delay was used to align microdialysis measurements with simultaneous vhFSCV recordings.

### Dopamine manipulation In Vivo

We used nomifensine (10mg/kg) and raclopride (1mg/kg), prepared in sterile PBS and administered intraperitoneally as pharmacological manipulation of striatal DA. The reward-expectation task is described in ^24^. Briefly, mice were water-restricted and, initially, habituated to lick a reward spout in response to randomly delivered rewards while head-fixed. After acclimation, they performed a task where rewards were delivered after a variable delay from the trial onset. Each trial began with a 4.3 kHz tone that increased frequency to indicate progress to reward. We used nine frequencies: 4.3 kHz, 6.2 kHz, 8.3 kHz, 10 kHz, 12.4 kHz, 14.1 kHz, 16 kHz, 18.4 kHz, and 20 kHz. The delay to receiving a reward was randomly drawn from a uniform distribution (4–8 s). At the end of each trial, the tone ceased, and the solenoid delivered 3μL of water to a spout in front of the mouse. Licking behavior was detected using capacitive touch sensors (AT42QT1010, Sparkfun). For optogenetic stimulation of DA cells, DAT-Cre^+^ mice received unilateral 0.5 µL injections of cre-dependent virus (AAV5.Syn.FLEX.ChrimsonR-tdTomato) into the midbrain (3.2AP, 0.8ML, 4.5DV). After three weeks to allow virus expression, an optic fiber was implanted into the midbrain under NAc FSCV guidance and cemented in a location that yielded the strongest evoked DA release. During experiments, awake animals received pulse trains of 625nm LED light at 60Hz and 60 pulses to closely mimic DA release caused by a reward. LED stimulations were delivered in bursts of various ITIs (every 10, 20, 30, 50, or 60 seconds) in 10-minute blocks with a 5-minute rest period between the blocks.

### Data Analysis

All analyses used custom routines in MATALB. The raw FSCV data consisted of a matrix of PxT dimension, where P represents voltage potentials for triangular ramp (−0.4V-1.3V) sampled in 1000 evenly spaced intervals. T is the product of recording duration in seconds and the sampling rate (10Hz). These raw data were imported directly into the workspace without any preprocessing. The DA oxidation current is defined at the potential where the peak current is present (as in Fig1B). This potential is slightly variable between electrodes but was always within a 0.55V-0.68V range (oxidation points 280-320). To facilitate the automatic detection of this location across electrodes, we used a peak detection function on samples with DA. We averaged the current magnitude within a 20-point range centered at the identified peak, typically 290-310 (0.58V-0.65V). This quantity is used as the DA oxidation current (^OX^DA), and used to calculate the ratio of these currents at the two scans (^OX^DA-Ratio). Specifically, ^OX^DA-Ratio is the ratio of ^OX^DA_(Scan-1)_ / ^OX^DA_(Scan-2)_. Note that the denominator is the scan with a smaller Hold duration. Serotonin and pH studies were performed using the same waveform as DA, and ^OX^5-HT was defined at 0.4V. Similarly, ^OX^pH was quantified at 0.1V.

All alignments used digital timestamps stored simultaneously with FSCV data. To generate calibration curves in Figures 3 and 4, we average peak oxidation currents across three analyte applications (as shown in Figure 2D) relative to the first data point and plotted current response under different analyte concentrations. Similarly, the Current-Ratio (^OX^DA-Ratio, ^OX^5-HT-Ratio, or ^OX^pH-Ratio) was first evaluated for the full-session data, aligned and averaged within and across electrodes under various analyte concentrations and Hold durations. Voltammogram correlations (Figure 3E) were performed by first averaging ^OX^DA_(Scan-1)_ and ^OX^DA-Ratio voltammograms for repeated analyte administration and compared using Pearson’s correlation.

Analysis of *in vivo* vhFSCV followed the same procedures described above for *in vitro* data processing. Simultaneous vhFSCV and microdialysis sessions were synchronized by initiating their acquisitions simultaneously. We performed spectral decomposition of wideband vhFSCV using the Matlab *spectrogram* function with logarithmically spaced frequencies from 1/sample duration to Nyquist frequency of data acquisition. Phasic transients quantified in Figure 6B and D were detected on a 0.5Hz highpass filtered data with the *findpeaks* function. We included detected peaks with a minimum transient width (at half max) of 250ms. To assess if phasic transients predict DA tone fluctuations, we counted the number of phasic events within a 30-second analysis window to determine the transient rate and evaluated the mean amplitudes of these transients. These transient frequency and amplitude quantities were used as predictors in a regression analysis. This tonic level is evaluated by lowpass filtering the broadband vhFSCV signal with a 0.03Hz cutoff frequency and average in the same 30s bins. We then lagged the phasic frequency/amplitude predictors relative to the tonic quantities in the 30-second bins to facilitate regression analysis of past phasic events predicting future DA tone levels. We performed the linear mixed effect regression using the *fitlme* function in Matlab using the following expression: TonicDopamine ∼ frequencyPhasicTransients * exptCondition + amplitudePhasicTransients * exptCondition + interactionTerm * exptCondition + (1 | mouseID). Finally, the same 30s-binned dataset, without lag, was used to determine their lead-lag relationships using the *xcorr* function.

